# BIFROST: a method for registering diverse imaging datasets of the *Drosophila* brain

**DOI:** 10.1101/2023.06.09.544408

**Authors:** Bella E. Brezovec, Andrew B. Berger, Yukun A. Hao, Albert Lin, Osama M. Ahmed, Diego A. Pacheco, Stephan Y. Thiberge, Mala Murthy, Thomas R. Clandinin

## Abstract

The heterogeneity of brain imaging methods in neuroscience provides rich data that cannot be captured by a single technique, and our interpretations benefit from approaches that enable easy comparison both within and across different data types. For example, comparing brain-wide neural dynamics across experiments and aligning such data to anatomical resources, such as gene expression patterns or connectomes, requires precise alignment to a common set of anatomical coordinates. However, this is challenging because registering *in vivo* functional imaging data to *ex vivo* reference atlases requires accommodating differences in imaging modality, microscope specification, and sample preparation. We overcome these challenges in *Drosophila* by building an *in vivo* reference atlas from multiphoton-imaged brains, called the Functional Drosophila Atlas (FDA). We then develop a two-step pipeline, BrIdge For Registering Over Statistical Templates (BIFROST), for transforming neural imaging data into this common space and for importing *ex vivo* resources such as connectomes. Using genetically labeled cell types as ground truth, we demonstrate registration with a precision of less than 10 microns. Overall, BIFROST provides a pipeline for registering functional imaging datasets in the fly, both within and across experiments.

**Significance:** Large-scale functional imaging experiments in *Drosophila* have given us new insights into neural activity in various sensory and behavioral contexts. However, precisely registering volumetric images from different studies has proven challenging, limiting quantitative comparisons of data across experiments. Here, we address this limitation by developing BIFROST, a registration pipeline robust to differences across experimental setups and datasets. We benchmark this pipeline by genetically labeling cell types in the fly brain and demonstrate sub-10 micron registration precision, both across specimens and across laboratories. We further demonstrate accurate registration between *in-vivo* brain volumes and ultrastructural connectomes, enabling direct structure-function comparisons in future experiments.

## Main

Calcium imaging studies of neural activity have provided central insights into brain function in multiple model systems, including the nematode *C. elegans* [1–6] the larval zebrafish [7– 11], the fruit fly [12–18] and the mouse [19]. In order to compare such volumetric imaging datasets across individual animals, data from individual animals is often aligned within a common set of spatial coordinates defining an atlas, an approach that has been widely used in fish, rodents and humans [20–23].

In this approach, the precision with which data can be registered to such a “local atlas” places limits on the effective spatial resolution of aggregated data, defining the spatial scale of quantitative comparisons. As it has proven challenging to precisely register data from different experiments in the same space, these atlases have generally been restricted to the bounds of a single project, where data was acquired using the same experimental apparatus and protocol [24–26].

The adult fruit fly *Drosophila melanogaster* is a well-established platform for circuits neuroscience and recent advances have enabled large-scale functional imaging in this system [12, 15–18, 27]. Such studies have revealed widespread sensory responses and movement-related neural activity, probed the relationships between neural activity and metabolism, and have led to the discovery of novel circuits. Each of these studies registered volumetric neural activity data either onto an *in vivo* local atlas or an extant *ex vivo* fixed-tissue atlas [28–35] However, different *in vivo* datasets have not been cross-registered, precluding direct comparisons, as well as a wealth of *ex vivo* neuroanatomical datasets [29], including gene expression patterns [36–38] and synapse-level wiring diagrams (connectomes) [32, 39–41] Cross-registration of these *ex vivo* resources has enhanced their utility as, for example, spatial registration has allowed morphologically defined cell types identified in the connectome to be associated with specific genetic driver lines [33, 35, 42, 43]. However, it has been difficult to align *in vivo* functional data to *ex vivo* atlases with cell-type precision (∼ 5*µ*m) [12, 15] due to the markedly different image statistics inherent to *in vivo* microscopy and fixed tissue imaging using light and electron microscopy.

Here, we present a robust and generalizable image registration pipeline, BrIdge For Registering Over Statistical Templates (BIFROST), that enables quantitative comparisons in *Drosophila*, across individuals and experimental setups. We created an *in vivo* atlas, the Functional *Drosophila* Atlas (FDA), that can accommodate functional datasets from different experiments and labs. An *in vivo* atlas allows us to represent functional activity in a common space which better reflects the geometry of the brain inside the head. We then aligned the FDA with extant *ex vivo* templates [28–31, 33–35], thereby importing atlas labels [30], neuropil annotations [30], information from the connectomes [32, 39–41], and powerful tools for neuron identification [33, 35, 42, 43]. Using these atlas labels, we demonstrate that our registration pipeline outperforms existing methods [44, 45]. We further validate our method by registering *in vivo* volumes collected on different microscopes in which the same cell types are fluorescently labeled to the FDA. Comparing these datasets in FDA space, we demonstrate that our cross-lab registration is precise to 5 microns. We also demonstrate that BIFROST can be used to align partial sub-volumes of the brain into FDA space, allowing users the flexibility to image particular regions of interest while retaining the ability to align to the atlas. Finally, we show that our pipeline can be used to register functional imaging data to connectomes with a precision of 5 microns. Thus, BIFROST creates a common space for *in vivo* neural imaging data, provides easy-to-use tools for accurate registration, and enables direct comparisons of functional data and *ex vivo* anatomical resources.

## Results

### Overview

Functional imaging datasets collected using fluorescence microscopy often comprise two separate channels, with one channel recording neuronal activity using one sensor (such as a calcium indicator), and one channel recording signals associated with a structural marker that broadly labels the brain. In our approach, the structural signals from individual brains in a single experiment are first registered together to form a template. The warp parameters derived from this transformation are then applied to the neuronal activity channel from each brain, thereby bringing these signals into the template space. Next, templates derived from each experiment or laboratory are aligned to the Functional Drosophila Atlas (FDA), allowing all datasets to be quantitatively compared to each other, and to other resources that are registered to the FDA.

### Developing the Functional Drosophila Atlas

Our goal was to develop an accurate pipeline for registering brain-wide imaging data to a single atlas. In flies, previous work has described atlases that span the entire brain using *ex vivo* datasets, and as well as atlases that span the central brain *in vivo* [12, 15, 29, 35]. However, no *in vivo* atlas spanning the entire brain has been described in either sex. To develop an atlas that best captures the structure of the female fly brain *in vivo*, a widely used model, we sought to suppress both individual and technical variation. To do this, we first imaged each individual brain, inside the head of the living fly, 100 times at a resolution of 0.6 × 0.6 × 1 µm, capturing expression of a pan-neuronally expressed cell surface marker (myristylated tdTomato) using two photon microscopy. These 100 volumes were then aligned using linear (affine) and non-linear (Symmetric Normalization (SyN)) transformations, as implemented in Advanced Normalization Tools (ANTs) [44, 45]. These were then averaged to define a single volumetric image of each brain that suppressed technical variation in each collected volume. This process was repeated for 30 individuals, and based on a qualitative assessment, 16 were selected for further image processing. Each of these images were normalized, sharpened, and iteratively aligned using linear and non-linear transformations to construct the FDA (Fig. S1A, Fig. S2, see Methods).

We next tried to align *ex vivo* resources, including JRC2018F anatomical labels and genetic tools, the hemibrain connectome and the FlyWire Connectome to the FDA [35, 39, 41]. This is a challenging registration problem because the image statistics associated with these imaging modalities have substantial differences that reflect (1) changes in brain morphology due to physical constraints of the head, (2) distortion created by fixation, and changes in the angle of the imaging axis (3) differences in the spatial distribution of fluorescence signals due to *in vivo* labeling of cell membranes versus *ex-vivo* immunohistochemistical labeling of synaptic antigens and (4) differences in SNR characteristics associated with single and two-photon microscopy. We initially attempted this alignment using ANTs; however, many regions of the brain aligned poorly.Therefore, to improve the registration, we adapted SynthMorph, a learned contrast-invariant registration method, and used it in sequence with linear and non-linear SyN transformations to improve registration of the *ex vivo* resources to the FDA [44, 46].

### Registering individual datasets to FDA

We collected neural activity (nSyb>GCaMP6s; the dependent channel) and anatomical data (nSyb>myr::tdTomato; the alignment channel) at brain wide scale in different labs using different imaging systems (Fig. 1, Methods). To register these datasets to the FDA, we first generated a dataset template by iteratively aligning the anatomical scan from each animal using linear and non-linear transformations (Fig. S1B, see Methods). We next used the combination of linear, non-linear SyN, and SynthMorph to register these anatomical scans to the FDA. The transformations that best align each anatomical scan were then applied to the corresponding neural activity data, thereby registering the functional signals to the FDA (Fig. S1C, see Methods).

**Figure 1.**
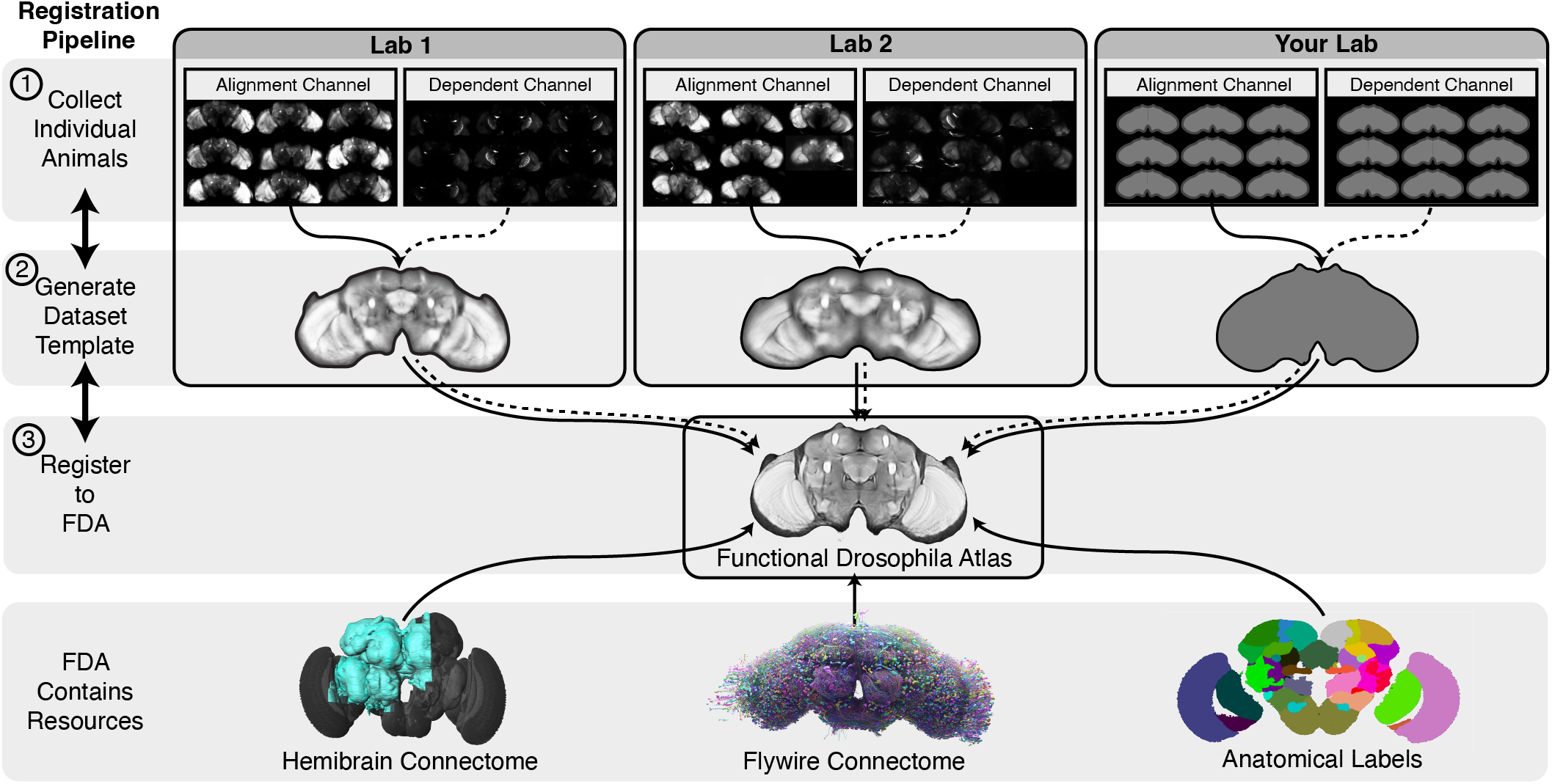
Overview of the BIFROST pipeline. (Step 1) Collect whole-brain volumetric data from multiple animals, with a pan-neuronal anatomical label used for alignment and an orthogonal dependent neural activity label. (Step 2) A dataset template is constructed, warping individual brains in the dataset to a common space. The template is constructed from the anatomical channels and the resulting transforms are applied to the neural data to register them into the template space. (Step 3) Dataset templates are aligned to the Functional *Drosophila* Atlas (FDA), in which all such datasets can be directly compared. Other resources have been registered to this space, including anatomical labels and connectomes.

### Quantifying registration performance

Making quantitative measurements of registration accuracy is challenging [47]. To address this challenge, we took two independent approaches. First, we quantified the performance of our method by measuring the overlap of small, well-defined anatomical regions that were manually labeled independently in both the *ex vivo* and *in vivo* atlases. Second, we expressed a fluorescent marker in cell-type specific sub-populations of neurons, and quantified their alignment within and across labs, and to connectomes.

### BIFROST outperforms existing methods for registration across modalities

We first quantified registration performance by measuring the alignment of neuropils labeled in the FDA space to the corresponding neuropils labeled in an established *ex vivo* anatomical atlas, JRC2018F (Fig. 2)[35]. Alignment accuracy was quantified for each pair of neuropils using the Sørenson-Dice coefficient, which captures the normalized fraction of voxels that overlap across both neuropil masks [48, 49]. For these analyses, we are calculating the transformation using the JRC2018F and FDA templates, and applying the transformations to the neuropil masks. As a control, we first used a linear transform to align the JRC2018F template to the FDA, and achieved an average Sørenson-Dice score of 0.52 (range: 0.13 to 0.75). Next, we added a non-linear transformation step (SyN), the core nonlinear transformation embedded in the widely used registration pipeline ANTs. However, SyN achieved only a modest increase in performance, with an average Sørenson-Dice score of 0.54 (range: 0.18 to 0.77), emphasizing the challenge of crossmodal registration. However, by adding SynthMorph to complete the BIFROST pipeline and perform the same alignment, we achieved an average Sørenson-Dice score of 0.65 (range: 0.45 to 0.84). We note that precision of registration did not deteriorate with tissue depth (Fig. S3). Thus, BIFROST provides an effective tool for registering signals across the brain.

**Figure 2.**
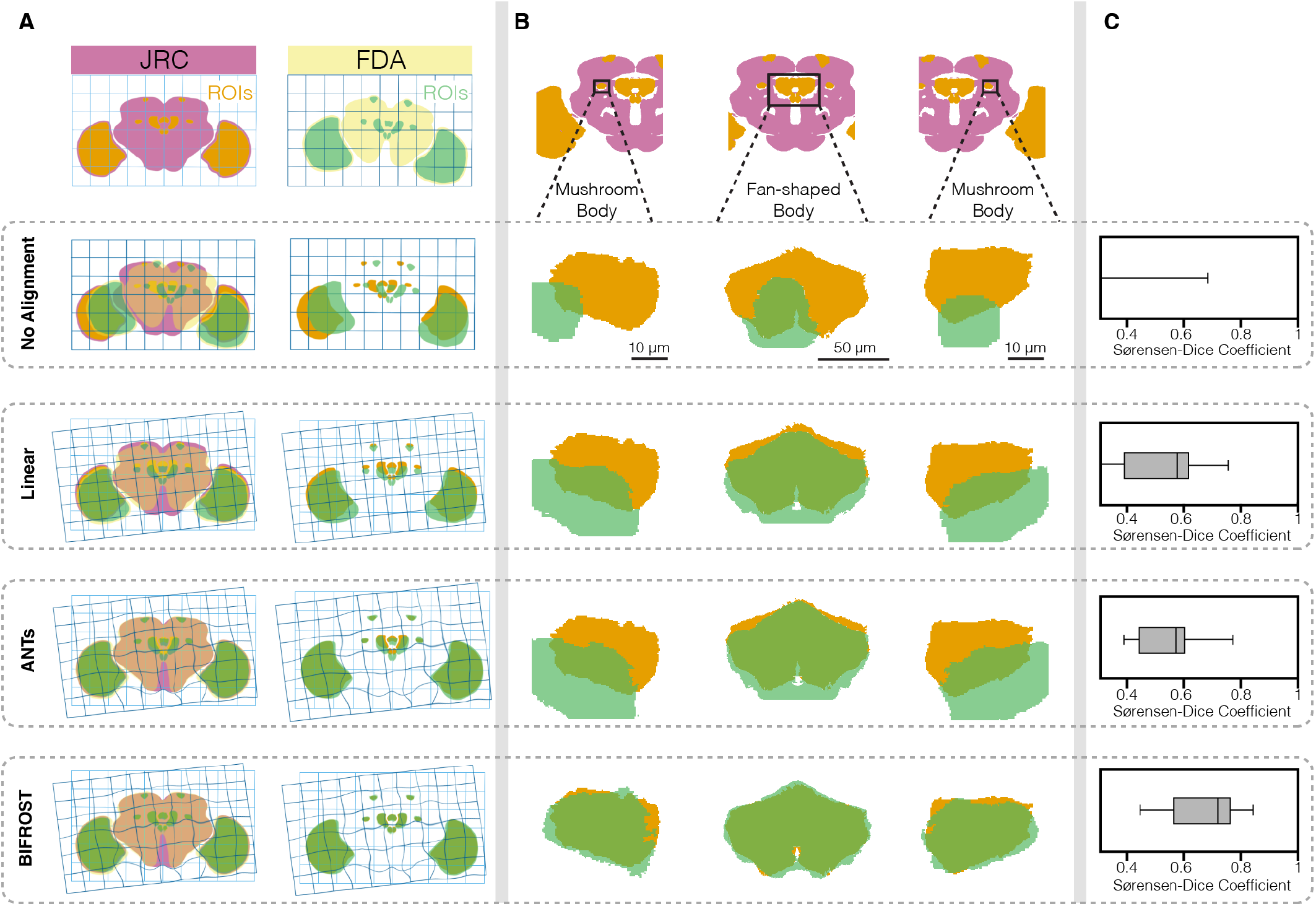
BIFROST registers neuropils across the brain with high precision. (A) Schematization of the registration steps of the BIFROST pipeline. (left) The FDA was transformed into the space of JRC2018F through successive applications of one linear and two non-linear SyN and SynthMorph steps. (right) The transformations computed at each step were applied to ROI annotations. Only a subset of the 13 labeled ROIs are illustrated. (B) Selected neuropil boundaries at successive steps of the pipeline. (C) Quantification of neuropil boundary overlap using the Sørenson-Dice coefficient. Box center line indicates median over all neuropils, box limits indicate quartiles, whiskers indicate minimum and maximum.

### Quantifying registration accuracy using sparse cell populations

While the Sørenson-Dice coefficient of labeled anatomical ROIs is widely used to estimate the precision of registration, this approach also has limitations [47]. The stereotyped architecture of the fly brain, combined with cell-type specific genetic labelling, make possible a quantitative assessment of registration precision, giving access to ground truth measurements that are generally not possible in other experimental systems. We first expressed a fluorescent indicator in a single genetically-identifiable cell type, Lobula Columnar 11 neurons (LC11). We chose the LC11 population because LC11 axons converge onto a single glomerulus, facilitating precise estimation of glomerulus position in 3D (Fig. 3 and Fig. S4). This glomerulus lies in the posterior ventral lateral protocerebrum (PVLP) and posterior lateral protocerebrum (PLP), two large neuropils that displayed relatively low contrast in the structural imaging channel. Thus, aligning LC11 within and across laboratories provides a challenging test-case for the BIFROST pipeline. As above, we compared the performance of the BIFROST pipeline to alternative, truncated pipelines that omitted various alignment steps, and included images collected independently in two laboratories (Fig. 3B). Each image was from the same strain, and expressed the neural activity marker GCaMP6s only in LC11 (as the dependent channel), as well as myristylated-td-Tomato in all neurons (as the structural channel). Qualitatively, individual LC11 glomeruli from both laboratories were similar in appearance after registration (Fig. 3C).

**Figure 3.**
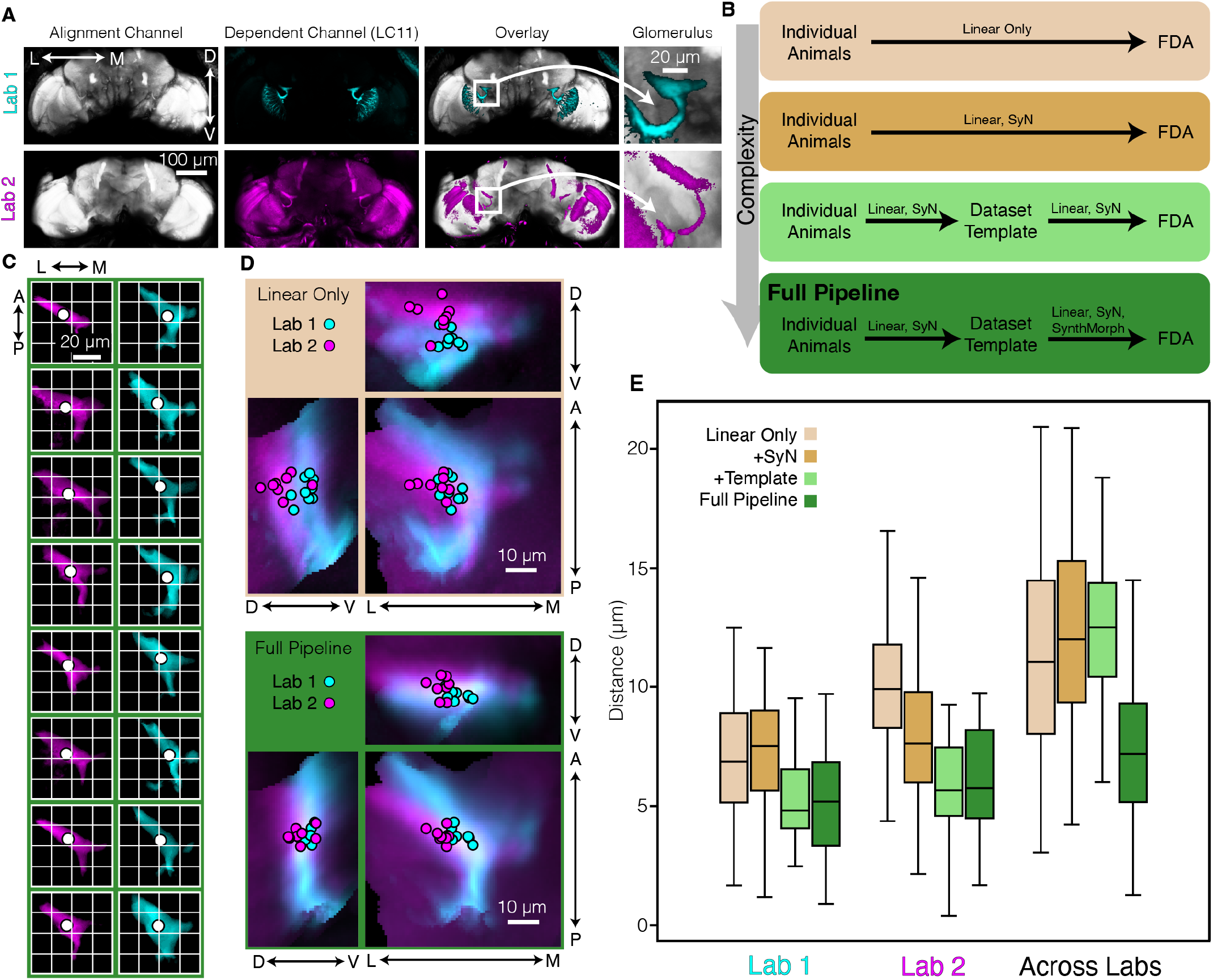
BIFROST registers brains within and across laboratories. (A) A single brain from each laboratory expressing an alignment channel (pan-neuronally expressed myr-tdtomato) and dependent channel (LC11-expressed GcAMP6s). The LC11 glomerulus region is highlighted. (B) Schematic flow chart of the BIFROST pipeline, as well as truncated versions that omit individual steps. (C) High magnification views of LC11 glomeruli in individual animals from both laboratories after registration into the FDA using BIFROST. Dot denotes centroid of each glomerulus. (D), As in (C), but individual glomeruli are overlaid and projections along each axis are shown. For comparison, glomeruli transformed by the linear-only pipeline and the full BIFROST pipeline are overlaid(Lab 1 n=9; Lab 2 n=8). (E) Quantification of the distribution of pairwise centroid distances within and across laboratories, for each pipeline variant. Box center line indicates median, box limits indicate quartiles, whiskers indicate 1.5x the inter-quartile range.

We quantified alignment precision by measuring the brainto-brain variation in the position of the centroid for each glomerulus, independently for both hemispheres, a feature that was robust and not strongly affected by threshold choice (Fig. S4A). The average pairwise displacement of any two centroids was 5.2 µm in Laboratory 1, 6.1 µm in Laboratory 2, and 7.3 µm across laboratories (Fig. 3D,E and Fig. S4B-D). We observed a nearly uniform error distribution, even including along the Z axis (corresponding to the anterior to posterior axis of the brain), the axis that generally suffers most from image distortion (Fig. S4E). Notably, the BIFROST pipeline outperformed all truncated variations of the pipeline, particularly for comparisons between labs (Fig. 3E and Fig. S4D). In addition to quantifying LC11 glomerulus registration using measurements of centroids, we manually labeled three additional glomerulus landmarks: the lateral tip, the medial tip, and the stalk. After registration, brain-to-brain variation in the position of the medial tip and the stalk was approximately equivalent to that seen with the centroids (Fig. S4F). However, brain-to-brain variation in the position of the lateral tip was increased across all samples, and across labs, all three landmarks were somewhat less precisely aligned than the centroid with an average pairwise displacement of 16.3 µm for the lateral tip, 10.3 µm for the medial tip, and 8.5 µm for the stalk (Fig. S4F). Some of this increased variation likely reflects the challenges of manual labeling; however, we also note that the lateral tip of LC11 glomerulus is relatively superficial and might be more affected by variation in the surgical preparation. Nonetheless, as discussed further below, the LC11 datasets derived from both labs aligned well with connectomic resources (Fig. 5).

To test whether this registration precision could be extended to a different brain region, we repeated this experiment using an additional cell population labelled by doublesex (DSX), a neuronal population that extends throughout many neuropils and comprises only fine <10 µm diameter) processes (Fig. S5A,B). Again, BIFROST achieved comparable results, displaying 5.5 µm average pairwise displacements between centroids in a readily identifiable structure within the DSX-expressing neuronal population (Fig. S5C,D).

### Registration of brain sub-volumes

Many experiments capture neural activity signals from only a sub-region of the brain and would benefit from registration across animals. We therefore adapted the BIFROST pipeline to align sub-volumes into the FDA (see Methods). To test the accuracy of sub-volume alignment, we generated a simulated subvolume dataset by selecting a 95 × 95 × 38 µm sub-region of one hemisphere from each LC11 brain (Fig. 4). Importantly, this sub-volume was not selected from the LC11 template; rather, it was selected independently for each fly, blind to variation in brain orientation and position. We then constructed a subvolume template from the individual sub-volumes (Fig. 4B,C). We aligned this sub-volume mean to the FDA using BIFROST (Fig. 4D,E) then assessed the accuracy of alignment as before (Fig. 4F-H). We found good agreement between the two sets of aligned data, with an average pairwise displacement between centroids of approximately 6 µm, demonstrating that the BIFROST pipeline can align partial brain volumes to the FDA.

**Figure 4.**
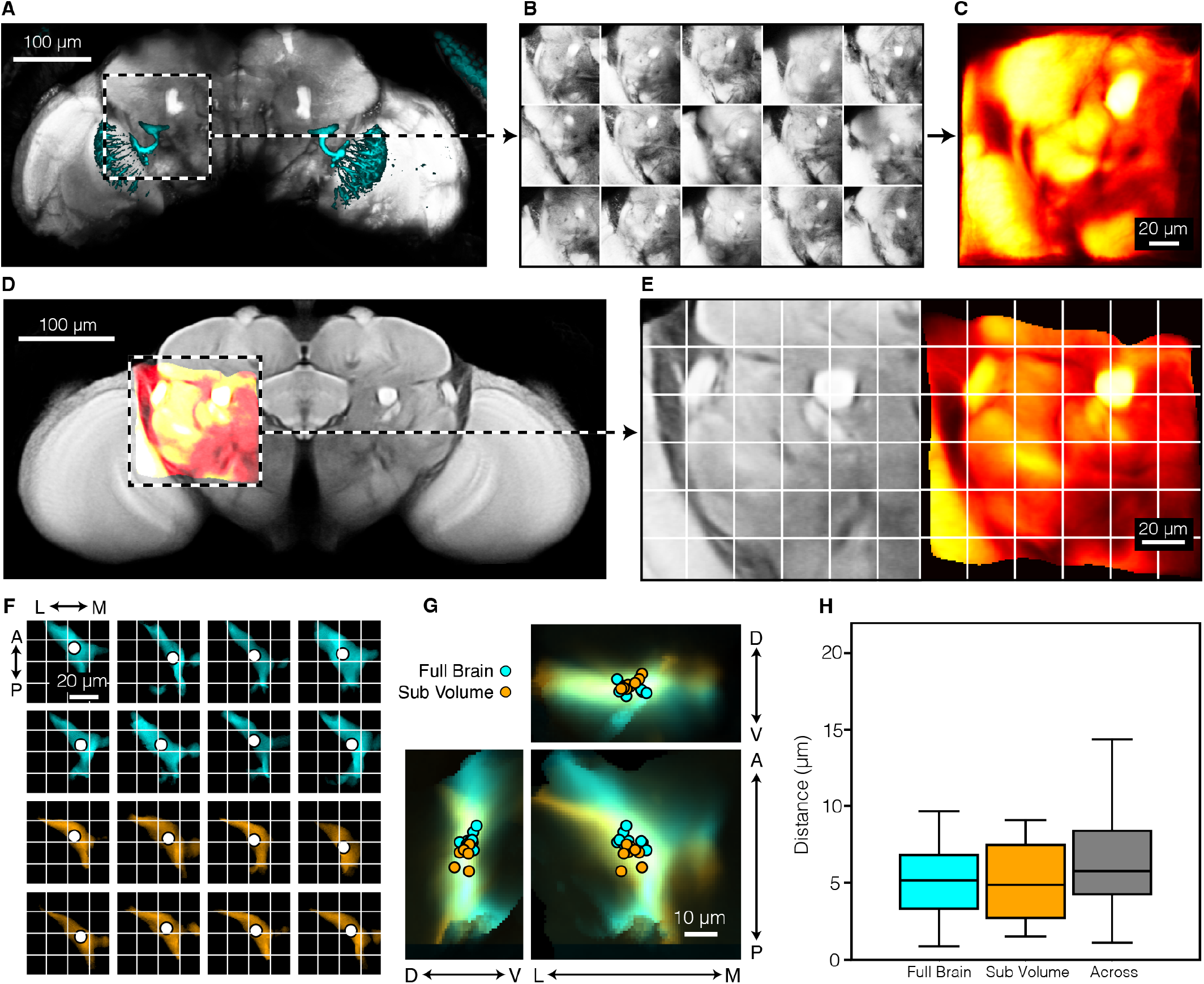
Sub-volumes of the brain can be accurately registered. (A) Example fly showing the alignment channel (pan-neuronally expressed myr-tdtomato, gray) and the dependent channel (LC11 expressed GcAMP6s, cyan). Dashed white box indicates LC11 glomerulus region. (B) Each brain sub-volume from each individual animal. (C) The dataset template generated from sub-volumes. (D) The template sub-volume (red) was aligned to the FDA (grey) using BIFROST. (E) High magnification view of the aligned mean sub-volume. FDA (left, grey) and aligned sub-volume (right, red) are shown. (F) High magnification view of each LC11 glomerulus after registration to FDA. Aligned LC11 glomeruli from either the whole brain (cyan) or subvolume imaging (orange). Dot denotes centroid of each glomerulus. (G) As in (F), but overlaid across animals and projected along each axis. (H) Quantification of pair-wise centroid distances after alignment using either the full brain image or the sub-volume. Box center line indicates median, box limits indicate quartiles, whiskers indicate 1.5x the inter-quartile range.

### Registration of the FDA with connectomes

Our goal was to align neuronal skeletons and synapse positions derived from connectomes to the FDA. Prior work has described the coordinate transformations from both the hemibrain connectome and the FlyWire connectome to JRC2018F [32, 41, 50]; therefore, we created a coordinate transformation from JRC2018F to FDA using BIFROST. This allows the coordinates of neuronal skeletons and synapses to be transformed to the FDA space through the path Hemibrain to JRC2018F to FDA, and FlyWire to JRC2018F to FDA. We note that SynthMorph does not provide methods for transforming the coordinates of a point cloud, which is required for a connectome. Therefore, after calculating the SynthMorph transformation, we recalculated it using ANTs (Fig. S1D, Methods).

We next examined the accuracy of this coordinate transformation by comparing the positions of the LC11 glomeruli measured in our *in vivo* datasets to that identified in both the hemibrain and FlyWire (Fig. 5). Remarkably, this cross-modal alignment was as precise as the alignment across *in vivo* datasets, with a precision of approximately 5 µm (Fig. 5C-F) for LC11, and 7 µm for DSX (Fig. S5D). Thus direct comparisons between anatomical wiring diagrams and functional volumetric images are now feasible with high precision.

**Figure 5.**
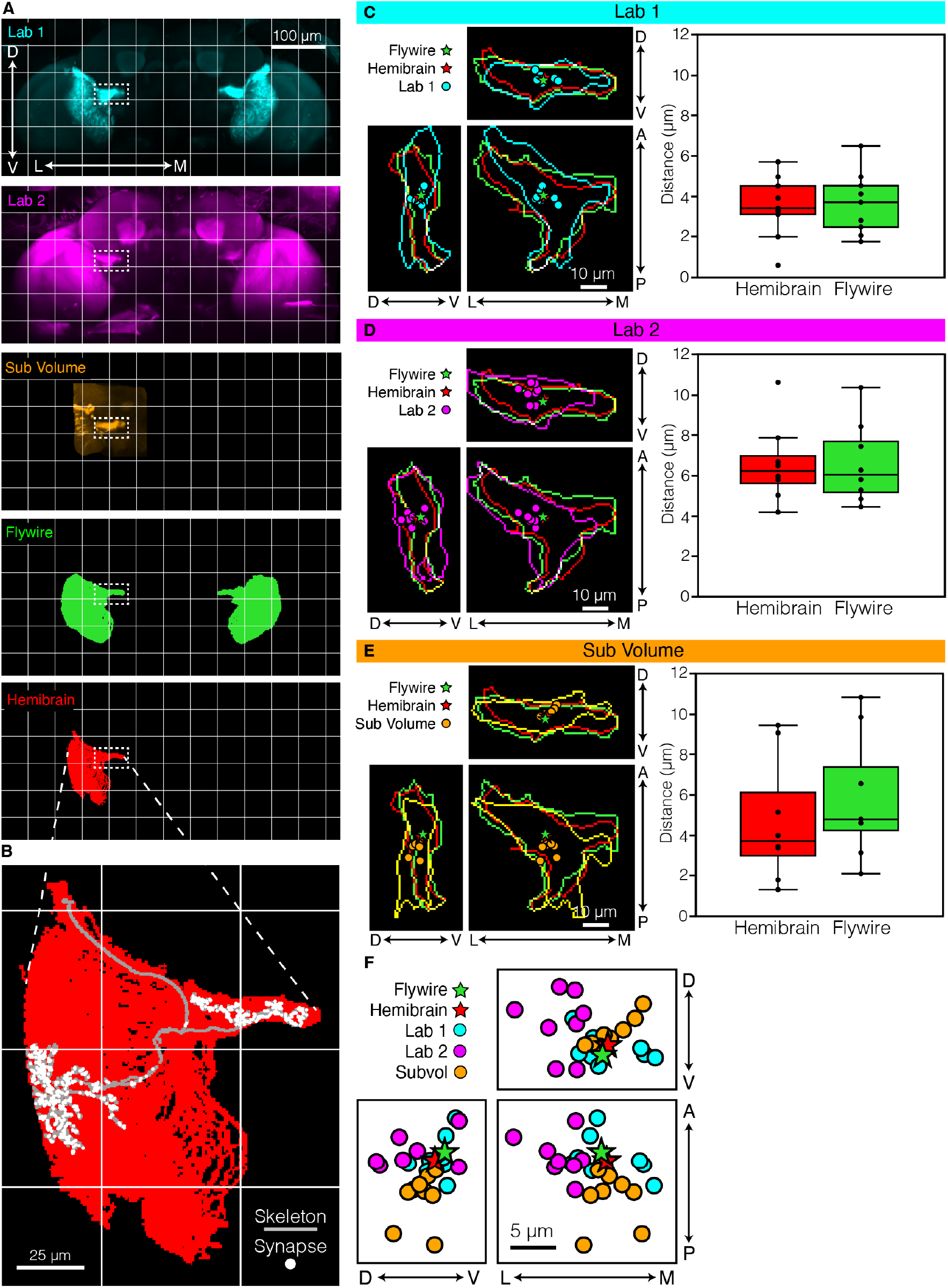
BIFROST enables registration of connectomes to *in vivo* imaging data. (A) Maximum projections of brains after alignment to the FDA. Template image of the LC11 channel for laboratory 1 (cyan), laboratory 2 (magenta), brain sub-volume (orange), LC11 skeletons from the flywire connectome (green), and LC11 skeletons from the hemibrain connectome (red), are shown. (B) Example of a single LC11 skeleton and synapses after being aligned to the FDA. (C) Comparing alignment accuracy of Laboratory 1 with the hemibrain and flywire connectomes. Left: high magnification view of the LC11 glomerulus; projections along each axis are shown. Cyan contour demarks the edges of the template image of LC11 from laboratory 1, while red and green denote the edge of LC11 skeletons from the hemibrain and flywire connectomes registered into the FDA. Right: quantification of the distribution of pair-wise centroid distances between each individual LC11 glomerulus and the hemibrain and flywire connectomes. Box center line indicates median, box limits indicate quartiles, whiskers indicate 1.5x the inter-quartile range. (D) As in (C), but for Laboratory 2. (E) As in (C), but using the sub-volume. (F) Overlay of LC11 centroids from all brains.

## Discussion

We developed BrIdge For Registering Over Statistical Templates (BIFROST), a pipeline for registering volumetric neural activity data across specimens and *in vivo* imaging systems. To complement BIFROST, we created the Functional Drosophila Atlas (FDA), an *in vivo* atlas that defines a common space for registering neural datasets. As an additional resource, we also provide the codebase needed to generate FDA-like templates. Using genetically-labeled neuron populations as ground truth, we show that BIFROST registers neural data across functional datasets at a scale of less than 10 microns, comparable to previous fixed-tissue registrations [51]. We further register connectomes, anatomical labels, and genetic resources to the FDA, thereby relating functional neural data to these resources. This toolkit addresses the longstanding challenge of precisely registering brain volumes across experiments, and will allow quantitative comparisons of neural activity in *Drosophila* across diverse datasets.

### Enabling comparisons across experiments

Comparing signals across experimental conditions and animals is critical for understanding large scale patterns of neural activity across genetic backgrounds, sensory contexts and behavioral states. In the fly, large-scale imaging experiments have uncovered brainwide activity patterns correlated with metabolic processes, sensory processing, locomotion, and feeding [12, 14– 17, 27, 52]. By facilitating statistical comparisons through precise cross-registration of these data types, BIFROST enhances quantitative comparisons of neural activity across conditions, an essential step toward building more comprehensive representations of brainwide dynamics.

### Enabling function-structure comparisons

By achieving precise registration between functional volumes and connectomes, BIFROST allows population-level activity signals to be associated with particular candidate cell types. While previous work in the fly has compared function to structure at the level of brain regions (neuropils) [15, 53], the sub10 micron alignment demonstrated here will enable functionstructure comparisons at an unprecedented spatial scale. For example, this precision will enhance the interpretation of functional data by using the connectome to constrain local and mesoscale cell connectivity, putative neurotransmitters, and labeled cell types [54, 55]. Additionally, registering such data at brain scale will enable users to move between testing both small and large-scale circuit and computational models, including those that link neural activity to sensory input, internal states, and behavior.

### Limitations

Although the BIFROST pipeline is flexible with regard to the particular structural marker used, some method to record the anatomical structure of the brain is required. This will mostly likely mean that an imaging channel is reserved for structural measurements of the *in-vivo* brain using pan-neuronal expression of a fluorescent label, precluding the use of this channel for recording a functional signal.

BIFROST achieves a registration accuracy of less than 10 µm, which surpasses existing methods, but is still non-zero, constraining interpretation of registered data. For example, if *in-vivo* data is collected at single micron resolution, registration will reduce its spatial precision to the 7 µm accuracy of BIFROST. Therefore, any spatial structure that existed in the neural data at a spatial resolution of less than the precision of BIFROST will be blurred. This is particularly important when attempting to assign neural identities based on connectome alignment, where it will typically be impossible to assign a single functional voxel signal to a particular neuron. However, at the same time, if functional signals span multiple imaging voxels, these correlated voxels can be assigned to small populations of candidate neurons that can be functionally validated using other approaches [16]. Finally, additional strategies to limit or sparsen expression of the functional effector can likely be implemented in parallel with large scale imaging to facilitate single-neuron identification [56].

Finally, we note that the flies used to create FDA were collected from a particular imaging axis that falls between the anterior-posterior and dorsal-ventral axes. Given the pointspread-function of the excitation beam inherent in two-photonmicroscopy, this results in a slight reduction in imaging resolution along this axis, which is visible from the medial-lateral axis (Figure S2). Despite this imaging artifact, registration accuracy does not deteriorate along this axis (Figure S4E).

## Conclusions

BIFROST, together with the codebase for constructing functional atlases, can be adapted for future use in other model organisms. Large-scale functional imaging experiments, as well as whole-brain anatomical studies, are increasingly feasible in many systems, including worms, flies, fish, mice, and primates. Direct comparisons between such functional data and anatomical wiring diagrams have advanced our understanding of computation [3, 57]. Moreover, there is broad interest in using connectomic constraints to inform computational models of neural activity [53, 54, 58–6 Tools capable of bridging functional and anatomical imaging modalities via precise volumetric registration will enable finer structure-function comparisons.

## Methods

### Genotypes

Flies were grown at 25°C on molasses (Clandinin Lab) or cornmeal (Murthy Lab) media, and imaged at 3-5 days post eclosion. The flies used to generate the FDA were *w+/w+;UASmyr::tdTomato/UAS-GCaMP6f; nSyb-Gal4/+*. The flies used to label LC11 neurons were *w+/w-;nSyb-LexA,LexAopmyr::tdTomato/R22H02-p65ADZp;UAS-GCaMP6s/R20G06ZpGAL4DBD*. The flies used to label DSX neurons were *w+/w+;brp>STOP>v5-LexA,LexAop-myr::tdTomato/UASmyr::tdTomato;DSX-FLP,LexAop-GCaMP6s/nSyb-Gal4*.

### Mounting and Dissection - Clandinin Lab

Flies were immobilized using a chilled Peltier plate, then fitted into a mount comprising a 3D-printed plastic dish holding a steel shim to secure the head and thorax. To reveal the posterior surface of the head, the head was pitched forward around the medial-lateral axis by approximately 70° relative to the thorax. UV curable glue was applied to the dorsal part of the head, and on the dorsal thorax. A saline solution was added to the dish for dissection (103 mM NaCl, 3 mM KCl, 5 mMTES, 1 mM NaH2PO4, 4 mM MgCl2, 1.5 mM CaCl2, 10 mM trehalose, 10 mM glucose, 7 mM sucrose, and 26 mM NaHCO3). The posterior head cuticle was cut using a tungsten needle and removed to expose the whole brain. Dissection forceps were used to remove fat and trachea.

### Mounting and Dissection - Murthy Lab

Flies were chilled on ice and placed in a Peltier-cooled “sarcophagus” held at 4°C, with the head of the animal restrained in a 3D printed holder. We positioned the head at a 90° angle relative to the thorax and restrained it via UV-cured glue and wax. The holder was then filled with saline, and the cuticle on the posterior side of the head was removed using fine forceps (Dumont 5SF) and a sharp needle. Fat and trachea were removed before imaging.

### Two-Photon Imaging - Clandinin Lab

Imaging data was collected using a resonant scanning Bruker Ultima IV system with a piezo drive and a Leica 20x HCX APO 1.0 NA water immersion objective. Either a Chameleon Vision II femtosecond laser (Coherent), or a MaiTai BB (SpectraPhysics) was used to excite GCaMP and tdTomato at 920nm. A 525/50nm filter and a 595/50nm filter were applied to the GCaMP and tdTomato emission photons, respectively. Photons in both channels were collected simultaneously using two GaAsP photomultiplier tubes (Hamamatsu). 100 imaging volumes were collected at 0.6 × 0.6 × 1 µm (1024 × 512 × 241 XYZ voxels).

### Two-Photon Imaging - Murthy Lab

Imaging data was collected on a custom-built 2-photon resonant scanning microscope equipped with a Chameleon Ultra II Ti:sapphire laser (Coherent) and a 25x water immersion objective (Olympus XLPLN25XWMP2). Dissected flies were placed below the objective and perfused with saline. The laser was used to excite GCaMP and tdTomato at 920 nm, with a 520/70nm filter (Semrock) applied to the green channel and a 617/73nm filter (Semrock) applied to the red channel. Note, the slightly wider band-pass of this green filter (compared to Clandinin Lab) likely contributed to additional bleed-through of photons from tdTomato, as can be seen in Figure 3. Photons in both channels were simultaneously collected using GASP photomultiplier tubes (Hamamatsu). We recorded 100 whole-brain volumes at a resolution of 0.49 × 0.49 × 1 µm (1024 × 512 × 300 XYZ voxels), to a sample depth of 300 µm. The microscope was controlled by ScanImage.

### Creation of FDA

Each anatomical scan was created by first imaging the myrtdTomato signal 100 times at 0.6 × 0.6 × 1 µm (1024 × 512 × 241 XYZ voxels). These 100 volumes were averaged, then each volume was warped (linear and non-linear) to this mean using ANTs, thereby correcting for motion. These aligned volumes were then averaged, creating the anatomical scan for each brain. Scans were additionally processed with an intensity based masking (to remove any contaminating background signal outside of the brain), removal of non-contiguous blobs (to remove, for example, cuticle which is otherwise visible due to auto-fluorescence), and histogram equalization to brighten overly dark areas and darken overly-bright areas (which assists in allowing a more uniform registration, and not an overemphasis on simply the brightest regions). Each brain was mirrored across the Y axis, doubling our effective data to 32 brains. These 32 brains were all linearly aligned to a single seed brain chosen from the 32, and averaged (“linear0”). The 32 brains were linearly aligned to “linear0”, and again averaged, producing “linear1”. Next, the individual anatomical scans were sharpened using the scikit-image implementation of unsharp masking, and aligned again (linear and non-linear) to “linear”, and averaged to produced “SyN0”. The last step was repeated two more times to produce the final FDA. We found that sharpening the brains before “linear0” caused poor convergence of neuropil boundaries, while completely omitting it resulted in blurry neuropil boundaries.

### The BIFROST pipeline

The BIFROST pipeline comprises four steps. First, a dataset template is created from structural volumes from each animal. Second, this dataset template is registered to the FDA. Third, each timepoint from each dependent channel is registered to the dataset template. Finally, these registered data are transformed into FDA space using the transformation calculated on step two. We provide the BIFROST pipeline as a Snakemake workflow that describes the dependency structure of the whole pipeline [61]. This facilitates parallel execution of independent steps, and as a result BIFROST can be transparently scaled from local execution on a single machine to thousands of parallel jobs on a cluster. This parallelization is critical, because serial execution of the ANTs dependent steps over such large datasets would take weeks to months to complete. BIFROST can be executed on all common cluster scheduling systems including Slurm, PBS and SGE and on cloud services via Kubernetes and several common cloud APIs [61]. This implementation allows a dataset over any number of animals with any number of channels, each imaged for an arbitrary number of timepoints and stored following a particular directory structure to be quickly submitted to a cluster for parallel execution with only minimal customization.

### The BIFROST pipeline: creation of dataset templates

Dataset templates were constructed from structural volumes of each animal following a standard procedure [35]. These volumes were mirrored, doubling the effective sample size, and pre-processed with the scikit-image implementation of contrast limited adaptive histogram equalization (CLAHE) using a kernel size of 64 [62, 63]. Template construction begins with a single volume, chosen arbitrarily from the pre-processed volumes to serve as the initial template. Linear (affine) transformations then aligned each pre-processed image to this initial template. Next, the transformed volumes were averaged to obtain a new template. Following this linear iteration, template construction continued with several (typically four) iterations of non-linear alignment and averaging. In each iteration, individual images were non-linearly transformed to the current template using SyN. Next, the transformed volumes were averaged to obtain a mean volume and the transformations themselves were also averaged. To complete each iteration, the next template is obtained by transforming the mean volume through the inverse of the mean transformation [64, 65]. The fourth iteration of this cycle produces the final dataset template. The pipeline is outlined in Fig. S1B. We have released our tooling for template construction as part of our Python package.

### The BIFROST pipeline: registration with SynthMorph

We wrote a configurable tool for image registration which registers a “moving” image to a “fixed” image with successive linear, non-linear SyN and SynthMorph transforms. If necessary, both “moving” and “fixed” images can be downsampled to reduce memory burden and computation time. As we found that SynthMorph was essential to effective registration in the central brain, but not the optic lobes, we also added support for masking the SynthMorph transform and manually generated a mask for the optic lobes using Fiji/ImageJ [66]. Next, we pre-processed the “moving” and “fixed” images by re-scaling intensities to the interval [0, 1] and applied the scikit-image implementation of CLAHE [62, 63] with a kernel size of 64 and configurable clip limit. By default CLAHE was applied to the “moving” image but can be applied to the “fixed” image as well.

After this pre-processing, linear (affine) a nd non-linear (SyN) transforms registering the “moving” image to the “fixed” image were computed and applied in sequence using ANTs.

Next, a non-linear SynthMorph transform was computed. As SynthMorph is constrained by its architecture to a fixed inference volume of 160 × 160 × 192 voxels, images were first transposed to align the longest axis of the image with that of the inference volume and then downsampled to 160 × 160 × 192 voxels. Finally, SynthMorph inference was run on the downsampled images yielding a warp field.

Next, the warp field was transposed back to the original axis order and then optionally mirror symmetrized across an algorithmically selected mirror plane, obtained by searching for the plane that minimizes the root mean square distance between the “moving” image and its mirror. This was performed on the “moving” image after it was affine t ransformed t o t he “fixed” image. In our experience, when the, “fixed” image i s aligned such that the dorsal-ventral axis lies along a principal axis of voxel coordinates this procedure reliably recovers the intended dorsal-ventral/anterior-posterior mirror plane of the *Drosophila* brain.

Next, the (optional) mask was transformed through the linear and non-linear transformations. The warp field w as then up-sampled to the original size of the images and applied to the “moving” image at all locations outside the mask, yielding the final image. The pipeline is outlined in F ig. S1C. All transformations and meta-data needed to apply the full transform were saved in a HDF5 file [67]. All tooling for computing registrations and applying the resulting transforms are provided as part of our Python package.

### Manual segmentation of neuropils

We used ITK-SNAP to manually draw regions of interest in an early version of the FDA, and then registered into the final FDA [68, 69]. We used ITK-SNAP’s built-in contrast-based segmentation to delineate the boundaries of the whole brain. We then hand-segmented the mushroom bodies (calyx, peduncles, ventral lobes, and medial lobes), central complex (protocerebral bridge), and optic lobes in each z-slice of the volume.

### Calculation of Sørenson-Dice coefficients for the crossmodal quantification

Region of interest annotations in the space of JRC2018F were obtained by registering a previous template that was published with regional annotations into the space of JRC2018F [28]. Correspondences between these ROIs and those annotated in the FDA were identified m anually. G iven s ets *X* a nd *Y*, the Sørenson-Dice coefficient is defined as

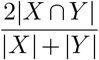

where |*X*| and |*Y* | are the cardinalities of the sets. The Sørenson-Dice coefficient was computed for each ROI in a voxelwise manner.

### Defining the positions of LC11 and DSX centroids

After alignment, whole-brain volumes were cropped to a region that contained the feature of interest (the terminal glomerulus for LC11, and a specific stalk for DSX). Box size was 95 × 57 × 76 µm for LC11, and 38 × 30 × 23 µm for DSX. In addition, for LC11, fluorescence outside of the PLP and PVLP regions were masked using the anatomical ROIs to avoid expression from LC11 dendrites in the lobula. An intensity threshold was then manually selected for each animal that best removed background fluorescence while maintaining the shape of the glomerulus. The image was then binarized and the center of mass determined

### Aligning JRC2018F and Connectomes to the FDA

The coordinates of the skeleton and synapses of LC11 were fetched from the online resources for the Hemibrain and FlyWire and were transformed into the space of JRC2018F using the flybrains Python package [70– In the flybrains package, the coordinate systems for the Hemibrain and FlyWire are labeled as “JRCFIB2018Fraw” and “FLYWIRE” respectively. These data were further transformed from JRC2018F into the space of the FDA by application of a bridging transformation, as follows. First, we applied the BIFROST pipeline to transform JRC2018F to the FDA (see Methods, The BIFROST pipeline: registration with SynthMorph). We next wished to apply this tranformation to the connectomes. However, SynthMorph does not support coordinate transformations, which is required for a connectome. Therefore, we recapitulated the full BIFROST transformation using only ANTs. We achieved this as follows. First, the FDA was transformed to JRC2018F using BIFROST. This FDA in JRC2018F space is now our new “fixed” target. Since this is now a single modality problem, we were then able to use ANTs to transform the original FDA to this fixed target. The pipeline is outlined in Fig. S1D.

## AUTHOR CONTRIBUTIONS

T.R.C., M.M., and B.E.B. conceived the project. B.E.B., Y.A.H., A.L., O.M.A., and D.A.P. collected data. B.E.B., A.B.B., Y.A.H., and O.M.A. analyzed the data. A.B.B. built the pipeline. S.Y.T. built the microscope used to collect data in the Murthy Lab. B.E.B., A.B.B., Y.A.H., A.L., O.M.A., and T.R.C wrote the manuscript. M.M. and T.R.C. advised throughout the project. B.E.B, A.B.B, Y.A.H, A.L. and O.M.A. contributed equally and have the right to list their name first in their CV.

## ACKNOWLEDGEMENTS

We would like to thank Megan Wang, Max Aragon, and members of the Clandinin lab for helpful discussions. We would also like to thank Greg Jefferis for sharing his advice, as well as Niyathi Annamaneni for help with figure design. OMA was supported by the Burroughs Wellcome Fund Postdoctoral Enrichment Program, the BRAINS Fellowship, the Simons Collaboration on the Global Brain Bridge to Independence Award, and the Princeton Presidential Postdoctoral Fellowship. BEB was supported by a National Science Foundation Graduate Research Fellowship. YAH was supported by a Stanford Bio-X Bowes Fellowship. AL was supported by the NSF through the Center for the Physics of Biological Function (PHY-1734030). This work was supported by NIH NINDS R35 to MM, R01EY022638 to TRC, NIH BRAIN R01 NS110060 to MM and TRC, Simons Collaboration of the Global Brain award to MM and TRC. TRC is a Chan-Zuckerberg BioHub Investigator.

## DATA AVAILABILITY

The FDA, bridging transforms and a replication dataset are available at Dryad: https://doi.org/10.5061/dryad.8pk0p2nx1

## CODE AVAILABILITY

Our software is available at Zenodo https://zenodo.org/doi/10.5281/zenodo.11097259

## Supplement

**Figure S1.**
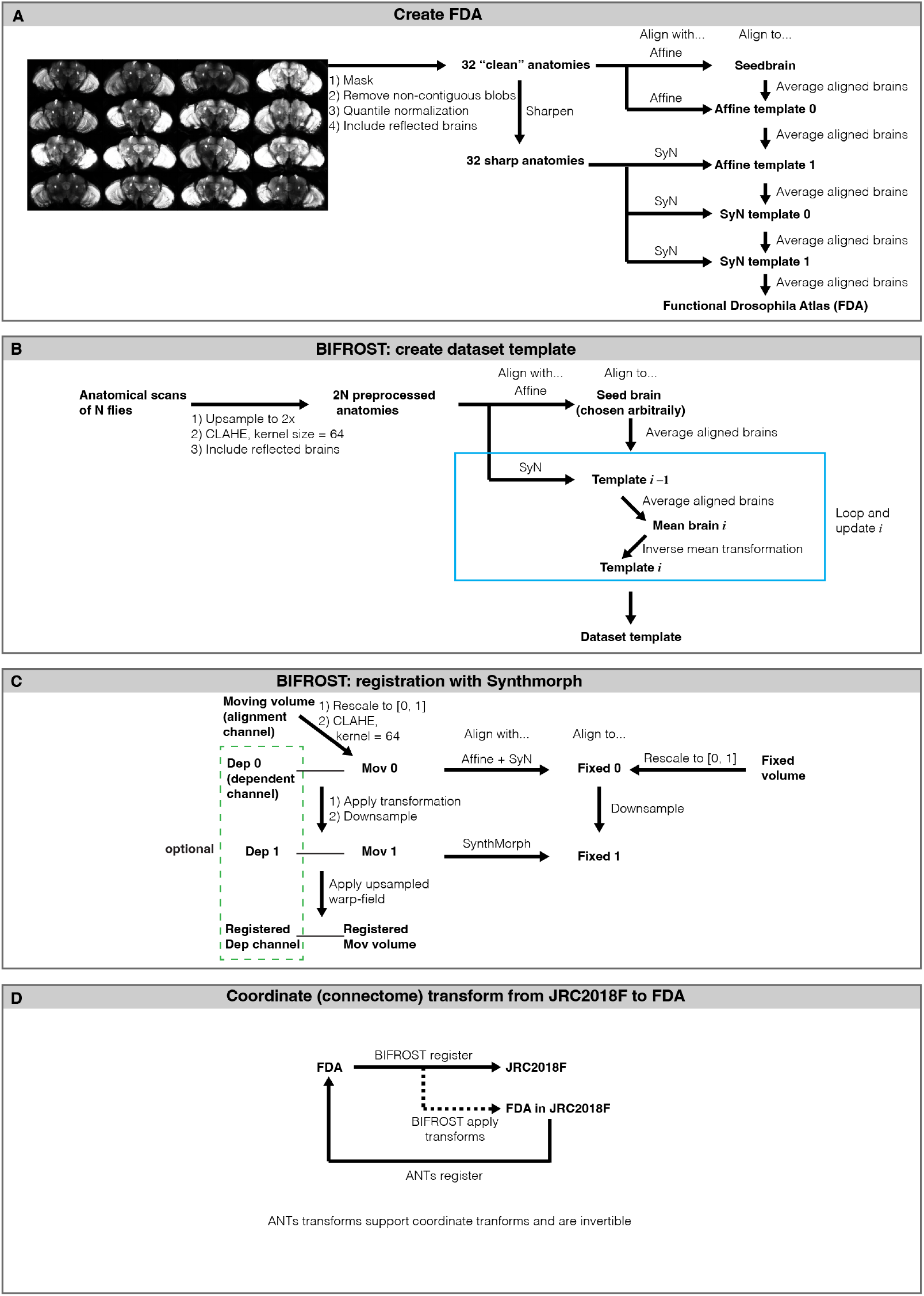
Creation of the Functional Drosophila Atlas (FDA) and BIFROST details. (A) Pipeline for creating FDA. (B) Pipeline for creating the dataset template. (C) Pipeline for registering a moving volume to a fixed volume. The alignment channel is used to register to the fixed volume, and the generated transformation is applied to the dependent channel, which is usually the functional imaging channel. (D) Pipeline for transforming the coordinates of a point cloud from JRC2018F to FDA. Note that ‘JRC2018F in FDA’ is JRC2018F registered to FDA using BIFROST pipeline outlined in (C). Full details in Methods.

**Figure S2.**
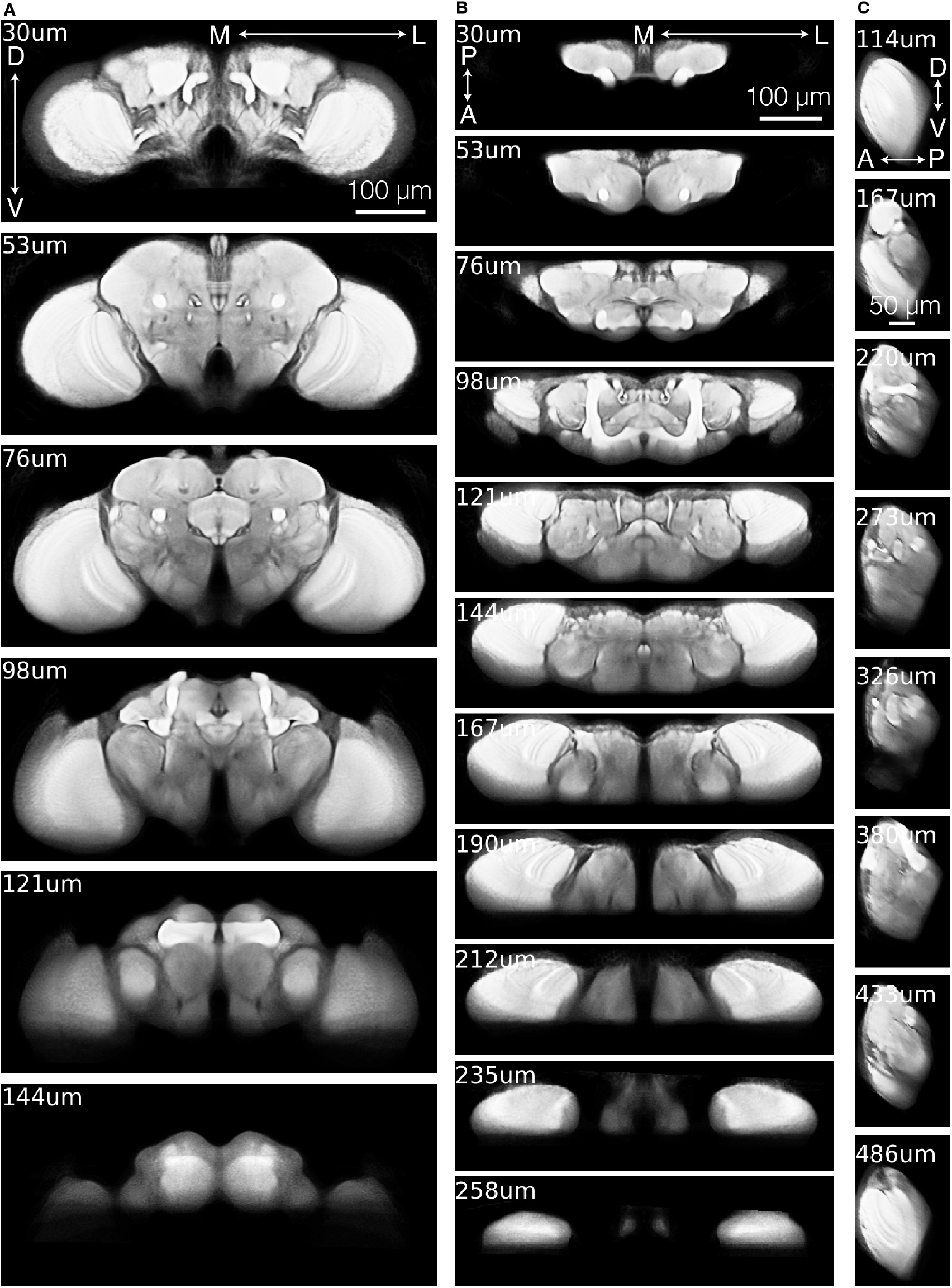
The Functional Drosophila Atlas (FDA). (A) Slices through FDA, moving along the anterior-posterior axis. Micron labels indicates depth along the axis. (B) Same as (A), except along the dorsal-ventral axis. (C), Same as (A), except along the medial-lateral axis.

**Figure S3.**
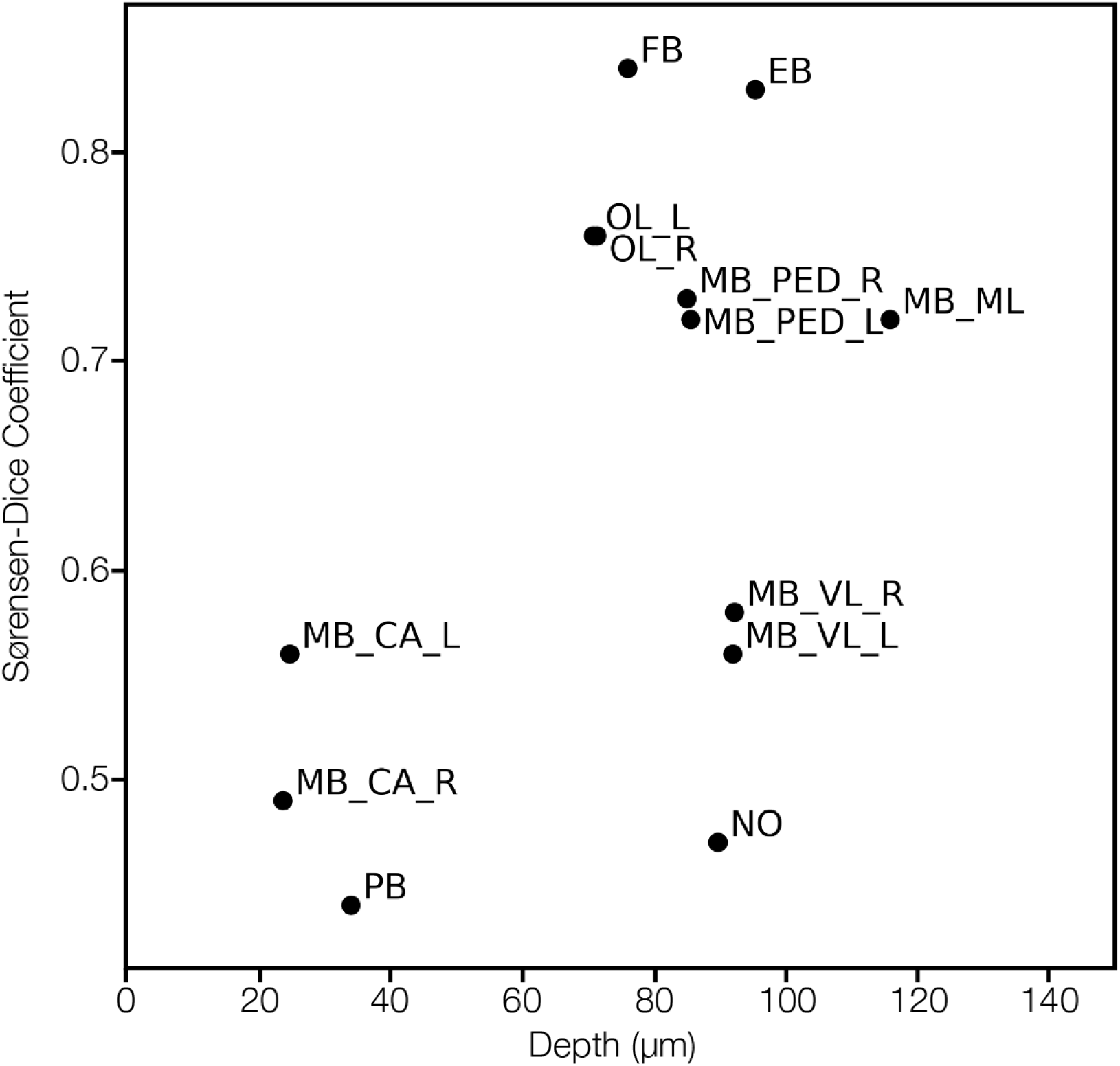
Registration accuracy does not deteriorate with tissue depth. Scatter plot related to Fig. 2 showing the Sørenson-Dice score of labeled neuropile versus their depth in the brain. Abbreviations: MB CA, Mushroom Body Calyx Left and Right; PB, Protocerebral Bridge; OL, Optic Lobes Left and Right; FB, Fan-Shaped Body; MB PED, Mushroom Body Peduncle; EB, Ellipsoid Body; MB VL, Mushroom Body Ventral Lobe Left and Right; NO, Nodulus; MB ML, Mushroom Body Medial Lobe.

**Figure S4.**
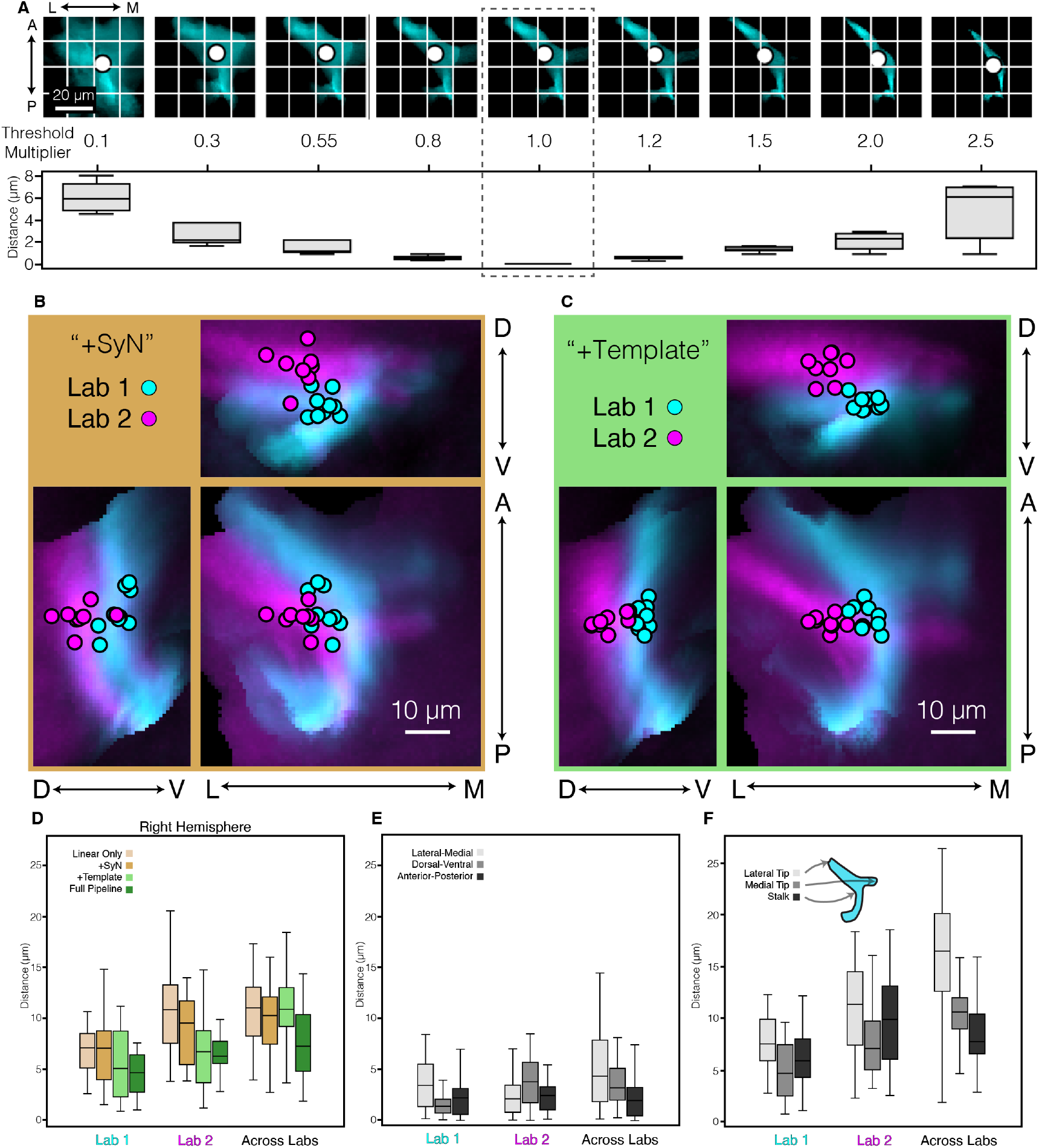
Quantifying LC11 registration accuracy. (A) Impact of threshold on centroid estimation. Throughout the manuscript, Otsu’s algorithm is used to select the threshold used for each animal. Here, we assess the impact of threshold value on centroid estimation by scaling the Otsu threshold by a threshold multiplier. For each animal, a range of threshold multipliers were applied, from 0.1 to 2.5. Top row, example LC11 glomerulus from a single animal as the threhold is adjusted. “1” is the original threshold. Bottom row, the 3D Euclidean distance between the original centroid and the adjusted centroid is calculated independently for each animal. Box plots show the distribution of centroid displacement from the original across animals. Box center line indicates median, box limits indicate quartiles, whiskers indicate 1.5x the inter-quartile range. Data from Lab 1 is used for this analysis. (B) Same as Fig. 3D, except showing the pipeline truncated to only ANTs linear and ANTs SyN (See Fig. 3B for schematic of truncations). (C) Same, except showing the pipeline truncated to only ANTs linear, ANTs SyN, and the creation of a Dataset Template (See Fig. 3B for schematic of truncations). (D) Same as Fig. 3E, except showing results from the other hemisphere. (E)Full pipeline, showing centroid distances along each orthogonal axis. Box center line indicates median, box limits indicate quartiles, whiskers indicate 1.5x the inter-quartile range. (F) Quantification of additional glomerulus landmarks. Instead of using the glomerulus centroid, three structures of the glomerulus are manually labeled for each animal: the lateral tip, the medial tip, and the point where the stalk meets the glomerulus. The distribution of pairwise distances is plotted for data aligned using the full pipeline.

**Figure S5.**
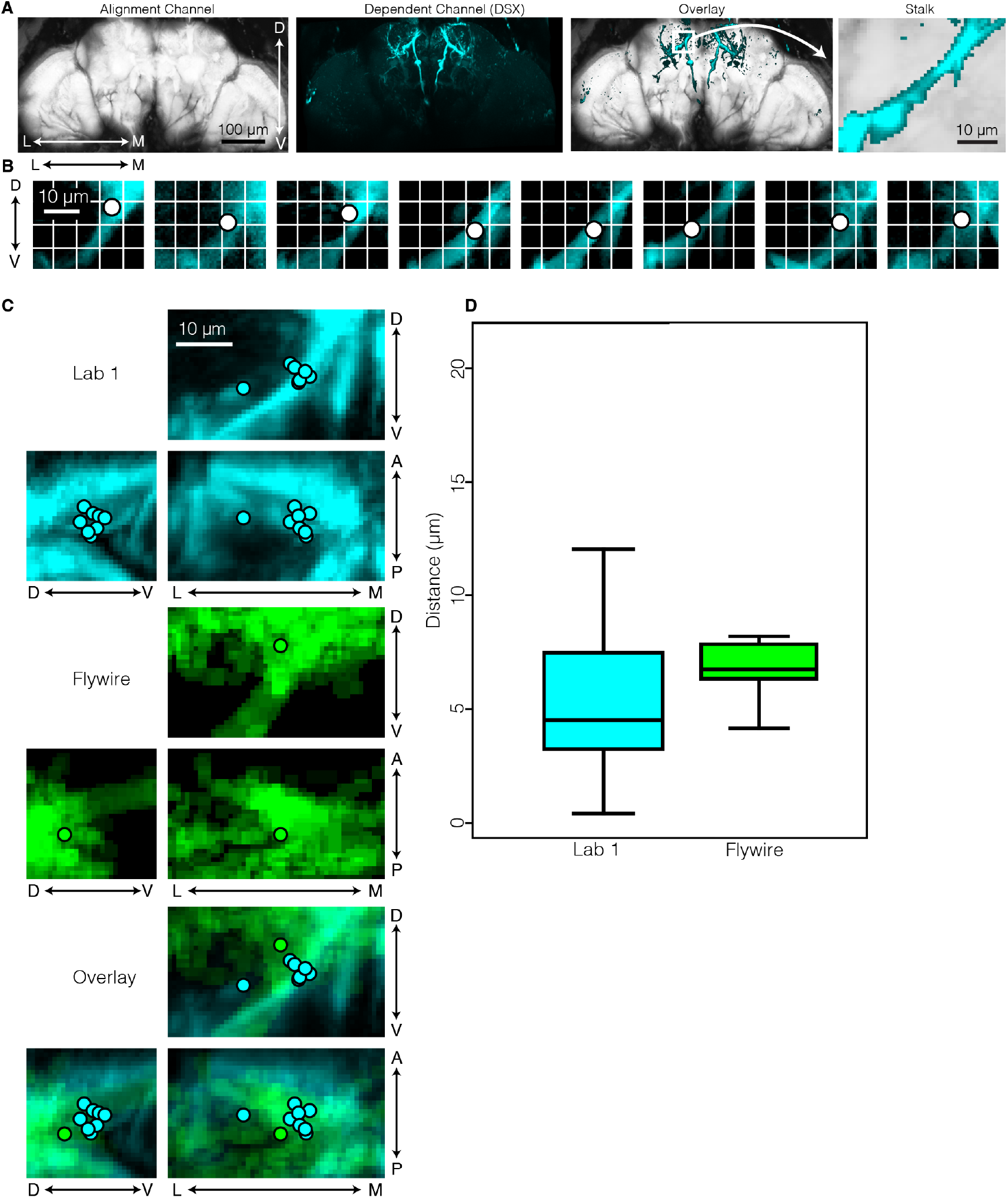
Quantifying DSX registration accuracy. (A) Example DSX fly showing the alignment channel (myr-tdtomato) and dependent channel (DSX). Zoom shows DSX stalk region that will be quantified. (B) Zoom in of DSX stalk region in FDA showing DSX expression of individual animals after the full pipeline was applied. Dot indicates centroid of each stalk, which will be used to quantify distance. (C) Same as in (B), but animals are overlayed and projections along each axis are shown. (D) Quantification of pair-wise centroid distances. Box center line indicates median, box limits indicate quartiles, whiskers indicate 1.5x the inter-quartile range.

